# Affinity Selection–Mass Spectrometry Coupled with Biophysical Validation Enables Proof-of-Concept Discovery of CHI3L1 Binders

**DOI:** 10.1101/2025.10.08.681114

**Authors:** Baljit Kaur, Hossam Nada, Moustafa Gabr

## Abstract

Chitinase-3–like protein 1 (CHI3L1) is a multifunctional extracellular glycoprotein implicated in tumor progression, immune suppression, and fibrosis, making it an attractive but challenging therapeutic target. To explore its chemical tractability, we applied an affinity selection–mass spectrometry (AS-MS) workflow to screen 10,000 small molecules for CHI3L1 binding. The screen yielded 124 initial hits with a hit rate of 1.24%, which were prioritized based on chemical suitability, and six candidates were advanced for validation using microscale thermophoresis (MST). Among these, compound **A9** exhibited a clear, dose-dependent binding response in MST with a Kd of 182 ± 18 µM. Molecular docking supported these findings, revealing that **A9** forms hydrophobic and hydrogen-bonding interactions within a defined pocket of the CHI3L1 structure. Although modest in affinity, **A9** represents the first small molecule binder of CHI3L1 identified through AS-MS. This study provides a proof-of-concept demonstration that CHI3L1 can be chemically engaged using AS-MS, establishing a foundation for future medicinal chemistry optimization and the development of chemical probes targeting this previously undruggable extracellular protein.

**Graphical Abstract:** 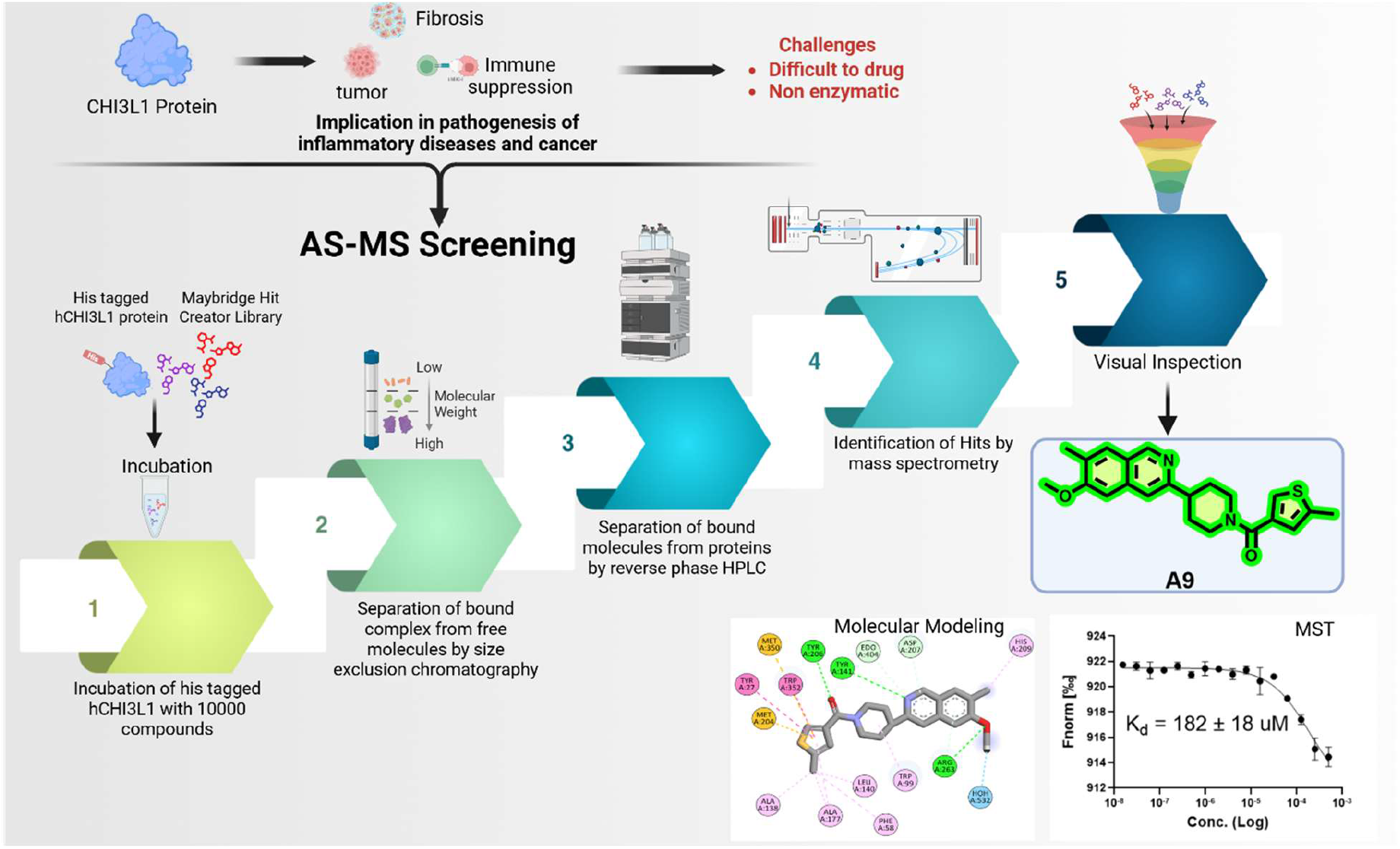

## 1. Introduction

The immune system relies on a network of extracellular proteins and signaling pathways to maintain homeostasis while mounting effective defense against pathogens and malignant cells.^1,2^ Dysregulation of these pathways can lead to chronic inflammation, fibrosis, or tumor progression.^3^ Among the many mediators influencing these processes, CHI3L1 (chitinase-3–like protein 1, also called YKL-40) is an extracellular glycoprotein with multiple functions, playing important roles in immune regulation, tissue remodeling, and cancer development.^4^ Multiple cell populations including macrophages, neutrophils, epithelial cells, fibroblasts, and tumor cells - secrete CHI3L1, enabling it to function across diverse tissue environments and contribute to a wide range of pathological conditions.^5^ Higher levels of CHI3L1 have been documented in multiple cancers, including glioblastoma,^6^ breast,^7–9^ colorectal,^10^ and non-small cell lung cancers,^11^ and are also associated with various inflammatory and fibrotic conditions, such as joint inflammation,^12^ respiratory disorders,^13^ and liver fibrosis.^14^ Clinical studies consistently associate high CHI3L1 levels with poor prognosis, aggressive disease, and increased risk of treatment failure, highlighting its relevance as a potential therapeutic target.

CHI3L1 mediates its biological effects through interactions with several receptors, including IL-13Rα2, syndecan-1, and integrins, which trigger intracellular signaling cascades such as MAPK/ERK, PI3K/AKT, and NF-κB.^15-19^ These pathways promote cell proliferation, survival, migration, and extracellular matrix remodeling. In the tumor microenvironment, CHI3L1 contributes to angiogenesis,^20^ supports metastatic spread,^21,22^ and suppresses antitumor immunity by polarizing macrophages toward an M2-like immunosuppressive phenotype^23-25^ and limiting T cell effector function.^26,27^ Similarly, in chronic inflammatory or fibrotic diseases, CHI3L1 drives persistent tissue remodeling, fibroblast proliferation, and excessive collagen deposition, exacerbating organ dysfunction.^28-30^ The ability of a single protein to influence such a wide range of cellular and tissue processes underscores its pleiotropic nature and central role in disease pathogenesis.^31,32^

Given its broad impact, CHI3L1 has emerged as an attractive but underexplored therapeutic target. Biologic inhibitors, such as monoclonal antibodies and soluble receptor constructs, have demonstrated preclinical efficacy in limiting CHI3L1-mediated tumor growth, inflammation, and fibrosis.^33,34^ However, these approaches face limitations in clinical translation, including restricted tissue penetration, immunogenic potential, and dependence on parenteral administration. Small molecules^35-37^ present a promising alternative due to their reversible, tunable, and potentially orally bioavailable properties. Yet, CHI3L1 has been difficult to target chemically because it is non-enzymatic and relies on flexible protein–protein interactions rather than well-defined catalytic pockets.

To explore whether CHI3L1 can be modulated with small molecules, we applied an affinity selection–mass spectrometry (AS-MS) strategy, which enables direct, label-free detection of protein–ligand interactions in solution. This approach is particularly well-suited for proteins like CHI3L1 that are challenging to assess using conventional activity-based screening. Screening a structurally diverse library led to the identification of 17 candidate compounds with potential CHI3L1-binding activity. These hits were further evaluated using orthogonal biophysical methods, including microscale thermophoresis (MST), to confirm direct engagement and measure binding affinities quantitatively.

This workflow led to the identification of a validated CHI3L1-binding compound. Although its affinity is in the high micromolar range, the ligand represents the first AS-MS–discovered small-molecule binder of CHI3L1. Our findings demonstrate that chemical matter can directly engage this challenging protein, complementing ongoing biologic strategies and establishing a tractable starting point for medicinal chemistry optimization. Beyond the specific ligand, this study highlights the utility of AS-MS as a discovery platform for extracellular, non-enzymatic targets such as CHI3L1.

## 2. Materials and Methods

### 2.1 Affinity selection mass spectrometry (ASMS) screening

#### Assay setup and conditions

ASMS was done using the Automated Ligand Identification System (ALIS) on an Agilent 2D-HPLC system interfaced to an Agilent Time-of-Flight MS. The assay conditions were 0.1 mM CHI3L1 using PBS (300 mM NaCl, 0.01% Tween20). Source of CHI3L1 for ASMS: human CHI3L1 Protein, His Tag (SinoBiological).

#### ASMS Method

The ALIS instrumentation consists of an Agilent 1260 HPLC pump for the sizeexclusion chromatography coupled to an Agilent 1290 UHPLC pump for the reversed phase (RP) chromatography with a high-pressure switching valve interfaced to an Agilent 6230B Time-of-Flight Mass Spectrometer. All MS acquisitions were done in positive ion acquisition mode.

Data analysis was performed with a combination of Agilent Mass Hunter and proprietary custom software.

- SEC conditions: Buffer A: 700 mM Ammonium Acetate Buffer B: 70% Acetonitrile Column: 50 × 2.1 mm Polyhydroxyethyl A, 3 mm 200 Å porosity (PolyLC)
- RP conditions: Buffer A: Water + 0.1% Formic Acid Buffer B: 90% Acetonitrile + 0.1% Formic Acid Column: 50 × 2.1 mm Kinetex 2.6 um C18 100 Å (Phenomenex).

Pools of compounds for a total of 10,000 compounds (from the Maybridge HitCreator and HitFinder libraries) at 10 μM were prepared in DMSO. Multiple copies of single use assay plates were prepared by dispensing 250 nL of each pool into a 384-well plate. The test compounds in DMSO were diluted to 10 μL in the assay buffer (PBS, 300 mM NaCl, 0.01% Tween20). One copy of the pools was run in a conventional HPLC-MS experiment to verify that the compounds could be detected. Data was analyzed using proprietary ASMS software. A total of 124 hits were identified.

### 2.2 Microscale Thermophoresis (MST)

#### 2.2.1 Single-Dose Screening

Protein–ligand interactions were assessed by MST using CHI3L1-His protein labeled with RED-tris-NTA fluorescent dye (NanoTemper Technologies) according to the manufacturer’s instructions. Labeling was performed by incubating 200 nM protein with 100 nM dye in 10 mM HEPES buffer (150 mM NaCl, 0.1% Pluronic F-127, pH 7.4) for 30 minutes at room temperature in the dark. The labeled protein was then diluted in PBS containing 0.05% Tween-20 and mixed with test compounds to achieve final concentrations of 20 nM protein and 250 μM compound. After incubation for 30 minutes at ambient temperature, samples were centrifuged briefly and analyzed on the Dianthus NT.23 Pico system. Buffer containing DMSO alone served as a negative control. All experiments were performed in triplicate, and mean values were reported.

To control for background fluorescence, test compounds (250 μM) were prepared from DMSO stocks in assay buffer, incubated in the dark, centrifuged, and analyzed on the same platform. Fluorescence quenching was assessed by incubating 10 nM dye with each compound under identical conditions.

#### 2.2.2 Dose–Response Analysis

Compounds identified as hits in the single-dose screen were further characterized in a dose– response MST assay. A 16-point serial dilution ranging from 500 μM to low nanomolar concentrations was prepared and mixed with labeled CHI3L1-His protein. The final DMSO concentration was adjusted to 4%. After 30 minutes of incubation at room temperature, samples were loaded into MST capillaries and analyzed on a Monolith NT.115 instrument using moderate-to-high IR-laser power and 60–80% LED intensity in the red channel. Binding affinities were determined using MO.Affinity Analysis software (NanoTemper).

### 2.3 Computational Study

Molecular docking studies were performed to investigate the binding mode of compound **A9** with CHI3L1 protein (PDB ID: 8R4X^37^) using the Schrödinger suite. The protein structure was prepared using the Protein Preparation Wizard, which included hydrogen addition, bond order assignment, and optimization of the hydrogen bonding network. The ligand structure was prepared using LigPrep, generating appropriate ionization states at physiological pH. Grid generation was centered on the co-crystalized ligand (XZ0) active site and docking was performed using Glide with standard precision mode. Molecular dynamics simulations were conducted using Desmond to assess the stability of the **A9**/CHI3L1 complex over 100 nanoseconds following previous protocol^36^. Briefly, the system was solvated in an orthorhombic TIP3P water box with appropriate counter ions to neutralize the system. Energy minimization was performed followed by equilibration using the default Desmond relaxation protocol. Production MD simulations were run at 300 K and 1 bar pressure using the NPT ensemble with periodic boundary conditions. Trajectory frames were saved at regular intervals for analysis. Root mean square deviation (RMSD) calculations were performed to evaluate the structural stability of both the unbound CHI3L1 and the **A9**/CHI3L1 complex throughout the simulation, and results were visualized using Xmgrace. Root mean square fluctuation (RMSF) analysis was conducted to identify residue-specific flexibility changes upon ligand binding. Protein-ligand contact analysis was performed to characterize the interaction patterns and calculate the interaction fraction for each residue throughout the simulation trajectory.

## 3. Results

### 3.1 Affinity selection–mass spectrometry (AS-MS) approach

To identify small-molecule ligands of CHI3L1, we employed an affinity selection–mass spectrometry (AS-MS) approach using our previously reported protocols^38-40^. We screened 10,000 compounds from the Thermo Scientific HitFinder library, which is designed to represent the chemical diversity of the full Maybridge Screening Collection. HitFinder compounds are selected using a clustering algorithm based on Daylight fingerprints and the Tanimoto similarity index (0.7 cutoff), ensuring broad coverage of drug-like chemical space. All compounds conform to Lipinski’s rules for “drug-likeness” (ClogP ≤5, hydrogen bond acceptors ≤10, hydrogen bond donors ≤5, molecular weight ≤500) and have a purity greater than 90%.

In the AS-MS workflow (Figure 1), CHI3L1 protein was incubated with compound mixtures to allow potential binders to form complexes. Protein - ligand complexes were separated from unbound molecules using size-exclusion chromatography (SEC), and bound compounds were subsequently released, resolved by HPLC, and identified by time-of-flight mass spectrometry. Initial screening across 10,000 compounds identified 124 potential hits with a hit rate of 1.24%. Manual filtering removed reactive, under-functionalized, and over-functionalized compounds, resulting in 17 candidates for validation (Figure 1).

**Figure 1.**
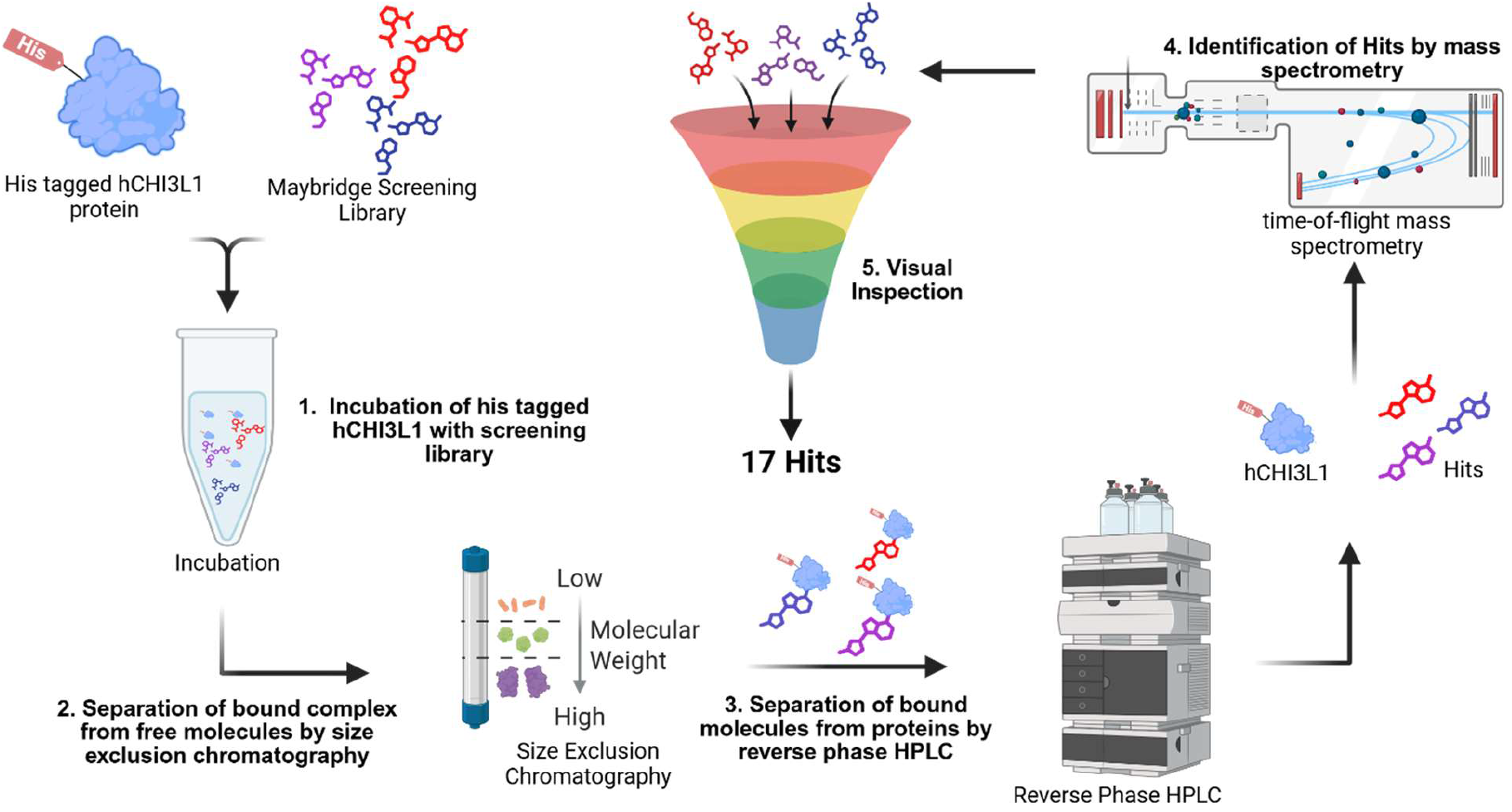
Schematic illustration of the ASMS-based screening workflow for identifying small-molecule candidates as CHI3L1 binders.

### 3.2 Primary screening by microscale thermophoresis (MST)

Seventeen candidate compounds were initially screened at a single concentration of 250 μM for binding to fluorescently labeled His-tagged hCHI3L1. Analysis of normalized fluorescence (Fnorm) values (Figure 2A) suggested 13 preliminary binders (**A1, A3, A4, A5, A6, A7, A8, A9, A10, B1, B3, B4**, and **B7**). To assess possible interference, compound fluorescence was compared with a reference containing labeled CHI3L1 and 2.5% DMSO in the absence of test compounds. This comparison revealed ten molecules (**A3, A4, A6, A7, A8, A9, A10, B1, B3**, and **B4**) with fluorescence intensities deviating significantly from the reference (Figure 2B).

**Figure 2.**
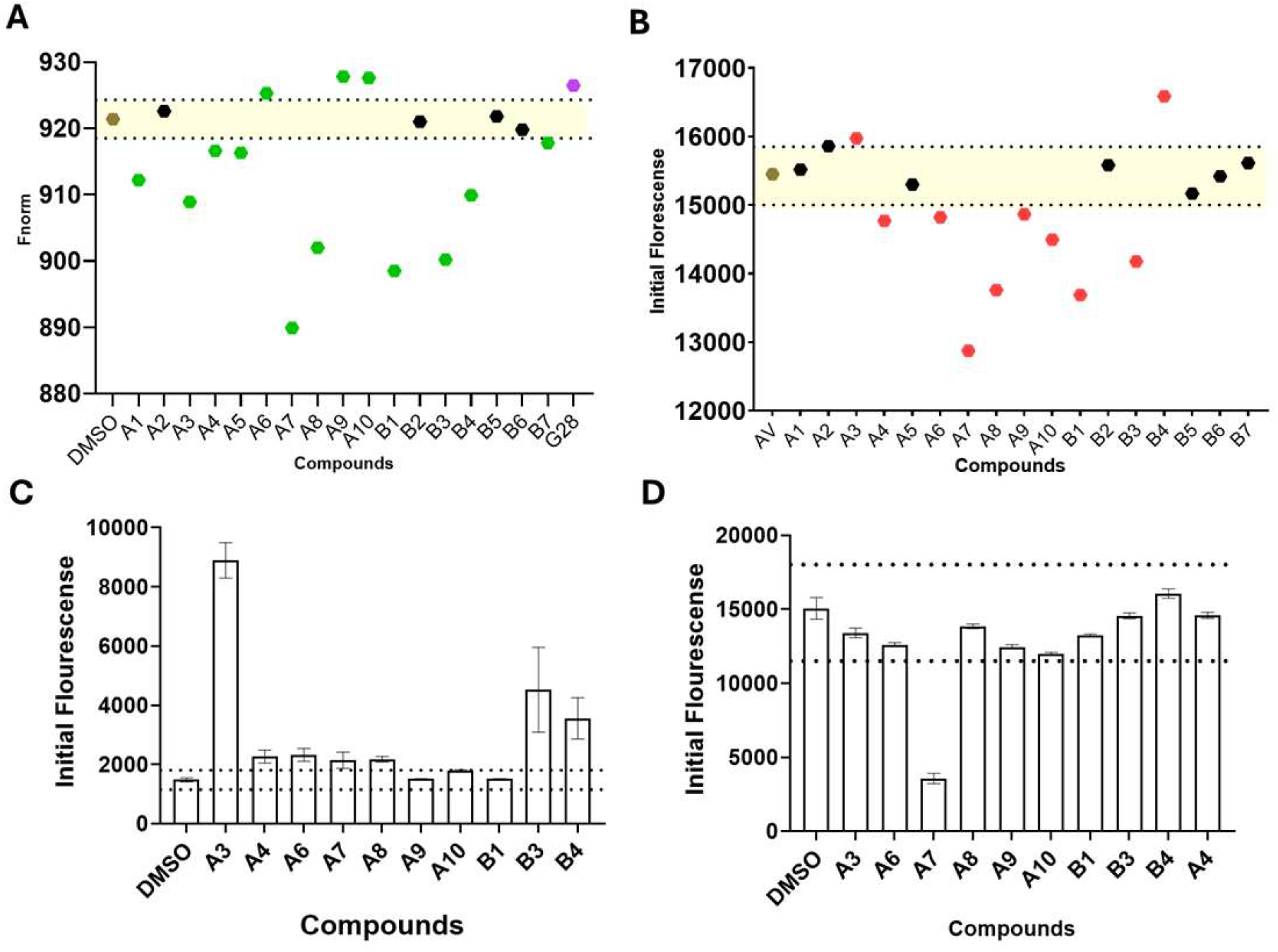
(A) Primary screening of compounds at 250 μM (with 2.5% DMSO). Green, brown, black, and purple indicate potential hits, negative control/reference (buffer with 2.5% DMSO), non-binders, and positive control (G28),^52^ respectively. (B) Comparison of initial fluorescence of compounds with the blank reference containing labeled hCHI3L1 and 2.5% DMSO. Red dots denote compounds that may exhibit signal interference or fluorophore interactions. **(C)** Autofluorescence and **(D)** quench test of candidates from single-dosage screening. Dotted lines in Figures 2C and 2D indicate the threshold values (mean negative control ± 5×SD) used to flag compounds with excessive autofluorescence or quenching.

To rule out false positives arising from intrinsic signal interference or dye interactions, additional controls were performed, including autofluorescence (Figure 2C) and quenching assays (Figure 2D). In the autofluorescence control, compounds **A3, A4, A6, A7, A8, B3**, and **B4** exhibited strong intrinsic fluorescence, whereas **A9, A10**, and **B1** remained within acceptable ranges for MST measurements. In the quenching assay, compound **A7** induced concentration-dependent quenching and was therefore excluded from further study.

Following this stepwise refinement, only compounds with minimal interference across all controls were considered reliable. Six molecules - **A1, A5, A9, A10, B1**, and **B7** - fulfilled these criteria and were validated as true hits with reproducible binding profiles.

### 3.3 Dose–response validation by Monolith

Compounds that passed the single-dose primary screen were subsequently evaluated in dose-response format using MST to confirm direct binding to CHI3L1 and to quantify binding affinities (Figure 3A). Fluorescently labeled His-tagged CHI3L1 protein was titrated with serial dilutions of each candidate compound over a concentration range of 1–1000 µM under optimized buffer conditions containing 2.5% DMSO. The resulting thermophoretic shifts were analyzed using the standard Hill model to derive dissociation constants (Kd). Of the six candidates, only compound **A9** (Figure 3B) demonstrated a clear concentration-dependent response with K_d_ of 182 18 μM, supporting its designation as a true binder. This moderate but measurable affinity typical of early-stage fragment-like ligands identified by AS-MS. The other candidates showed flat or noisy thermophoretic responses, suggesting either nonspecific association or insufficient binding strength under the tested conditions. Collectively, these MST data provide quantitative confirmation that compound **A9** interacts directly with CHI3L1 in solution, supporting its designation as a small molecule binder suitable for subsequent validation by molecular docking analysis.

**Figure 3.**
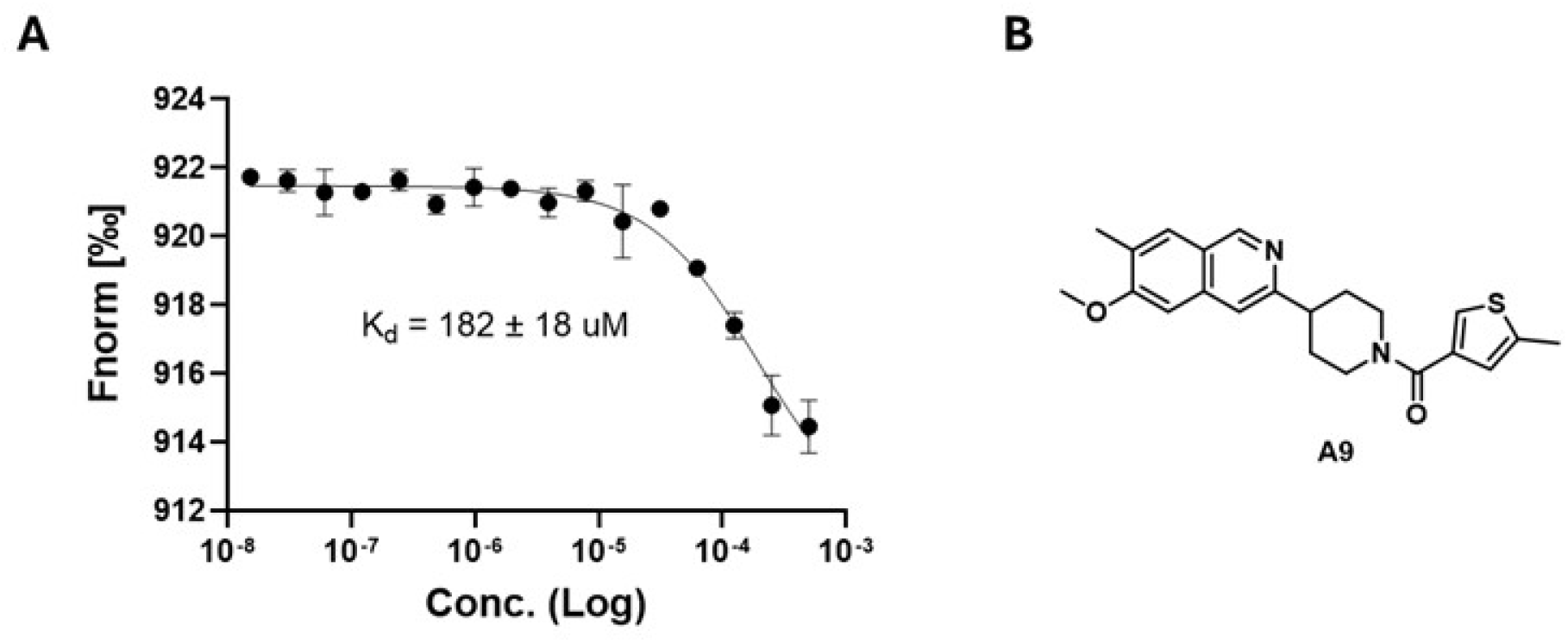
(A) Dose response confirmation of CHI3L1 binding by compound **A9** by MST. The graph displays dose-dependent changes in normalized fluorescence (Fnorm [%]) plotted against increasing concentration of ligand; (B) Chemical structure of **A9**.

### 3.4 Molecular Docking

To gain structural insight into the binding mode of compound **A9** and to rationalize the observed biophysical interactions, molecular docking and dynamics simulations were performed using the crystal structure of human CHI3L1. These in silico analyses aimed to identify potential binding pockets, key residue interactions, and the physicochemical basis for ligand recognition. The molecular docking and dynamics studies predicted that compound **A9** establishes interactions (Figure 4) with the CHI3L1 binding pocket through a network of hydrophobic, hydrogen bonding, and aromatic interactions. The 3D binding pose demonstrates that **A9** occupies a well-defined pocket within the CHI3L1 structure. The 2D interaction diagram (Figure 4C) reveals key binding interactions with hydrogen bonds established with Tyr141, Tyr206 and Arg263 amino acid residues of the binding site. The RMSD analysis (Figure 4D) over the 100 ns simulation period indicates that the **A9**/CHI3L1 complex maintains structural stability with RMSD values fluctuating around 1.2-1.4 Å for most of the trajectory which is comparable to the unbound CHI3L1 protein. These results suggest that ligand binding does not induce significant conformational changes or destabilization which conforms with the moderate binding activity observed experimentally.

**Figure 4.**
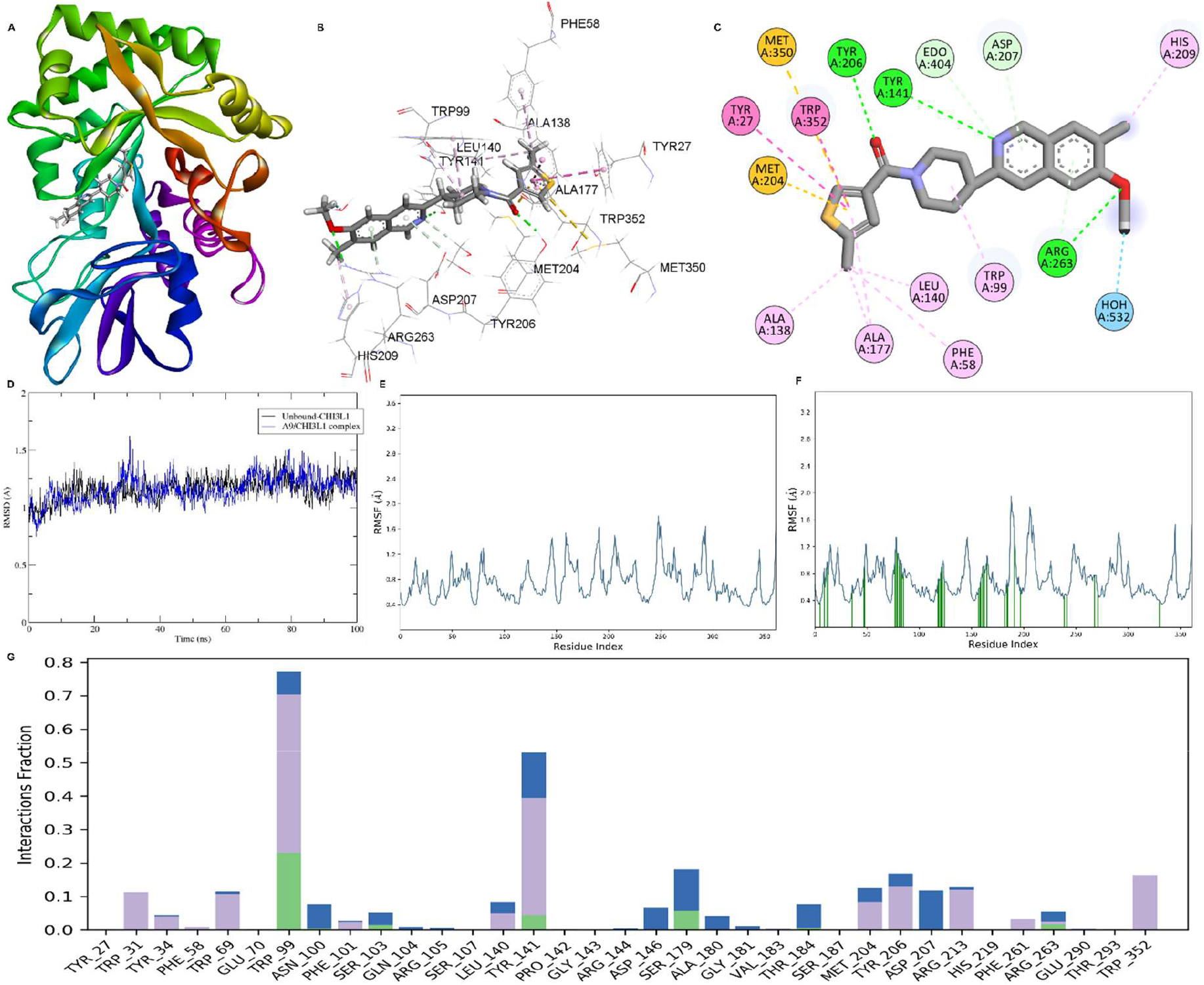
Molecular docking and dynamics simulation analysis of compound A9 binding to CHI3L1. (A) Three-dimensional representation of **A9** bound within the CHI3L1 protein structure (PDB: 8R4X). (B) Detailed three-dimensional view of the **A9**/CHI3L1 binding interface showing key interacting residues in stick representation. (C) Two-dimensional interaction diagram showing binding site residues represented as colored circles, with green indicating hydrophobic interactions, pink representing hydrogen bonds or polar interactions, orange showing ionic interactions, and dashed lines depicting interaction distances to the ligand (shown in stick representation). (D) RMSD plot of backbone atoms over 100 nanoseconds of molecular dynamics simulation, comparing unbound CHI3L1 (black line) and **A9**/CHI3L1 complex (blue line). RMSF profiles of the unbound protein (E) and **A9**/CHI3L1 complex (F). (H) Protein-ligand contact interaction fraction analysis.

The RMSF profiles (Figure 4E-F) reveal that both the unbound and bound forms exhibit similar flexibility patterns across most residues, with certain loop regions showing higher fluctuations. Notably, some regions display reduced flexibility upon A9 binding which suggests that the binding of **A9** to the CHI3L1 protein stabilizes these regions. The interaction fraction analysis (Figure 4G) demonstrates that Trp99 was predicted to establish the highest interaction frequency, followed by Tyr141, indicating these residues maintain persistent contacts with **A9** throughout the simulation. The stable RMSD profile, consistent RMSF patterns, and persistent protein-ligand contacts throughout the 100 ns simulation collectively indicate that **A9** forms a thermodynamically stable complex with CHI3L1, validating the predicted binding mode from molecular docking studies.

## 4 Conclusion

Targeting CHI3L1 with small molecules remains a challenging but promising strategy for modulating tumor growth, immune evasion, and fibrotic processes. Using AS-MS screening followed by orthogonal biophysical validation, we identified compound **A9** as a CHI3L1 binder with measurable though modest affinity. Rather than representing a final therapeutic agent, **A9** serves as a validated chemical starting point that demonstrates the feasibility of engaging CHI3L1 with small molecules. Future optimization and structural characterization will be essential to improve potency and selectivity, ultimately enabling the development of chemical probes and therapeutic candidates for cancer, fibrosis, and chronic inflammatory conditions.

## ACKNOWLEDGMENTS

We gratefully acknowledge financial support from the National Institute of Neurological Disorders and Stroke under grant number R01NS136524.

## Author Contributions

The manuscript was written through contributions of all authors. All authors have given approval to the final version of the manuscript.

