## Supporting Information for "Affinity Selection–Mass Spectrometry Coupled with Biophysical Validation Enables Proof-of-Concept Discovery of CHI3L1 Binders"

**Contents**

| **1.** | **Smiles of selected hits as potential CHI3L1 binders** | **2** |
| --- | --- | --- |

**Table S1. Smiles of selected hits as potential CHI3L1 binders.**

| **Codes** | **SMILES** |
| --- | --- |
| **A1** | CCc1c(snn1)C(=O)N2CCCC23CC(N(C3)c4cccc(c4)F)(C)C |
| **A2** | c1cc(ccc1CCC(=O)N2CCC(CC2)c3ccnc4n3ncc4)F |
| **A3** | c1ccc(cc1)CC(=O)NC2CCN(CC2)c3nc(cs3)c4ccccc4 |
| **A4** | COc1ccccc1/N=c2/c(cc3ccc(cc3o2)O)C(=O)NCC4CCCO4 |
| **A5** | Cc1c(c(n(n1)CC(=O)NCc2ccc(cc2)n3cccc3)C)[N+](=O)[O-] |
| **A6** | CC1(CC2(CCCN2C(=O)c3cccc(c3)COC)CN1c4cccc(c4)F)C |
| **A7** | Cc1c(c2ccccc2[nH]1)/C=C/c3nc4ccccc4c(=O)n3C |
| **A8** | CCOc1ccc(cn1)C(=O)N2CCCC23CC(N(C3)c4cccc(c4)F)(C)C |
| **A9** | Cc1cc2cnc(cc2cc1OC)C3CCN(CC3)C(=O)c4cc(sc4)C |
| **A10** | c1cc2c(cc([nH]2)C3CCN(CC3)C(=O)c4cc5cc(ccc5[nH]4)F)nc1 |
| **B1** | Cc1ccc(cc1)CNc2nc3c(n2CC(=C)C)c(=O)[nH]c(=O)n3C |
| **B2** | Cc1cc(nc(n1)NC(C)C(=O)NCC2(CC2)c3ccccc3Cl)C |
| **B3** | CC1(CC(c2ccccc2O1)NC(=O)Cc3csc(n3)c4ccccc4)C |
| **B4** | Cc1ccc(c(c1)CC(=O)N2CCC(CC2)(COc3ccc(c(c3)C)C)O)OC |
| **B5** | Cc1nc(cs1)CC(=O)N2C3CCC2Cn4c(nc5ccccc5c4=O)C3 |
| **B6** | c1cc(ccc1c2c[nH]nc2C3CCCNC3)C(F)(F)F |
| **B7** | CC1=C(C(C2=C(N1)CCCC2=O)c3ccccc3F)C(=O)N4CCOCC4 |
